# Transcription factor-like 5 is a potential DNA/RNA-binding protein essential for maintaining male fertility in mice

**DOI:** 10.1101/2021.06.17.448807

**Authors:** Weiya Xu, Yiyun Zhang, Dongdong Qin, Yiqian Gui, Shu Wang, Guihua Du, Fan Yang, Lufan Li, Shuiqiao Yuan, Mei Wang, Xin Wu

## Abstract

Tissue-specific transcription factors often play key roles in the development of specific cell lineages. Transcription factor-like 5 (TCFL5) is a testis-specific protein that contains the basic helix-loop-helix domain, although the *in vivo* functions of TCFL5 remain unknown. Herein, we generated CRISPR/Cas9-mediated knockout mice to dissect the function of TCFL5 in mouse testes. Surprisingly, we found that it was difficult to generate homozygous mice with the *Tcfl5* deletion since the heterozygous males (*Tcfl5*^+/−^) were infertile. We did, however, observe markedly abnormal phenotypes of spermatids and spermatozoa in the testes and epididymides of *Tcfl5*^+/−^ mice. Mechanistically, we demonstrated that TCFL5 transcriptionally regulated a set of genes participating in male germ cell development, which we uncovered via RNA-sequencing and TCFL5 ChIP-sequencing. We also found that TCFL5 interacted with RNA-binding proteins (RBPs) that regulated RNA processing, and further identified the fragile X mental retardation gene 1, autosomal homolog (FXR1, a known RBP) as an interacting partner of TCFL5 that may coordinate the transition and localization of TCFL5 in the nucleus. Collectively, we herein report for the first time that *Tcfl5* is haploinsufficient *in vivo* and hypothesize that TCFL5 may be a dual-function protein that mediates DNA and RNA to regulate spermatogenesis.

## INTRODUCTION

Infertility disorders affect approximately 15% of couples, and about half are caused by male factors(Schlegel, 2009)—with abnormal spermatogenesis the principal cause of male infertility(Silber, 1994). Spermatogenesis in mammals includes three primary stages: mitosis of spermatogonia, meiosis of spermatocytes, and spermiogenesis. Spermiogenesis is the last step of spermatogenesis in which the haploid round spermatids differentiate into mature spermatozoa amid a series of dramatic cellular reorganizations and morphologic changes—including acrosomal formation, nuclear elongation and condensation, flagellar development, and cytoplasmic elimination(Eddy, 2002). Precise genetic regulation during spermatogenesis is required for sperm production to take place in concert with functional integrity. Novel regulatory elements are therefore worthy of exploring since spermatogenesis represents one of the most complicated biologic processes *in vivo*.

Over the past several decades, key regulators—in particular the germline-specific factors in the process of spermatogenesis—have been reported to act via transcriptional and post-transcriptional regulation(Bettegowda and Wilkinson, 2010; Legrand and Hobbs, 2018). For example, testis-specific transcription factor CREMτ is a well-studied key transcription activator that controls post-meiotic germ cell differentiation. CREMτ is expressed in post-meiotic spermatids and directly targets a list of post-meiotic genes that encode structural proteins required for spermatid differentiation(Kosir et al., 2012; Martianov et al., 2010; Nantel et al., 1996). *Cremτ*-deficient mice exhibit complete arrest of spermiogenesis at step 5 during the 16-step formation process from round spermatids to elongated spermatids(Nantel et al., 1996). SOX family transcription factor SOX30 is a transcription factor revealed to control the gene-expression transition from late meiotic to post-meiotic stages, as well as subsequent development of round spermatids(Bai and Fu, 2018), and SOX30-mutant males are sterile owing to spermiogenic arrest at the early round-spermatid stage(Bai and Fu, 2018). In addition, post-transcriptional regulation of gene expression also plays critical roles in spermatogenesis. RANBP9 (Ran-binding protein 9) has been identified as a regulator of alternative splicing in spermatocytes and spermatids(Bao et al., 2014). In *Ranbp9-*knockout testes, expression of key spermiogenic genes is dysregulated and unique mRNA transcript isoforms are increased as a result of aberrant alternative splicing, leading to germ cell loss and infertility(Bao et al., 2014). TPAP (testis-specific poly(A) polymerase) has been demonstrated to maintain mRNA transcript stability(Kashiwabara et al., 2000) and is essential for the progression of haploid spermatid differentiation(Kashiwabara et al., 2002), and *Tpap*-null mice were infertile as a result of failed spermatid morphogenesis(Kashiwabara et al., 2002). SAM68 (Src-associated substrate in mitosis of 68 kDa) is important for alternative splicing regulation and translational control in male meiotic germ cells, with loss of *Sam68* resulting in aberrant differentiation of round spermatids, increased apoptosis, and consequent infertility(Paronetto et al., 2009). These post-transcriptionally regulated RNA-processing events such as pre-mRNA splicing, mRNA export, maintenance of transcript stability, and translation in the male germline are required to maintain coordinated gene expression(Legrand and Hobbs, 2018).

The basic helix–loop–helix (bHLH) family of transcription factors is receiving increased attention due to its critical roles during the development and differentiation of a wide variety of cell types (Davis et al., 1987; Massari and Murre, 2000; Murre et al., 1989). Our best-understood bHLH proteins are the myogenic regulatory factors (MRFs), which are capable of converting the mesodermal cell line C3H10T1/2 into muscle progenitor cells known as myoblasts(Olson, 1990). During spermatogenesis in the testes, studies have also revealed that multiple bHLH family members occupy essential functional roles in the development of germ cells. SPZ1 (bHLH-Zip transcriptional factor) is expressed in the testes and epididymides of adult male mice, and overexpression induces apoptosis of germ cells at an early stage of spermatogenesis, reduces the populations of normal spermatozoa, creates smaller litter sizes, and eventuates in infertility at 6 months of age (Hsu et al., 2004). Sohlh1 and Sohlh2 (spermatogenesis-and oogenesis-specific basic helix-loop-helix 1 and 2) are expressed in KIT-negative and-positive spermatogonia, and are both required for spermatogonial differentiation through the regulation of the expression of *Gfrα1*, *Sox3,* and *Kit* genes(Ballow et al., 2006a; Ballow et al., 2006b; Hao et al., 2008; Suzuki et al., 2012; Toyoda et al., 2009). SOHLH1 and SOHLH2 hetero-and homodimerize with each other *in vivo*, and double-mutants of *Sohlh1/Sohlh2* phenocopy the single mutant of *Sohlh1 or Sohlh2,* inducing improper spermatogonial differentiation and prematurely activating meiosis (Suzuki et al., 2012).

Transcription factor-like 5 (TCFL5) was previously considered to be a testis-specific protein, and the encoded domains and localization in the spermatocyte subsets of the testes are conserved between humans and mice(Maruyama et al., 1998; Siep et al., 2004); however, the *in vivo* function and mechanism(s) underlying TCFL5 action have not yet been elucidated. In the present study, we characterized the expression of TCFL5 in mouse testis, demonstrated that the loss of TCFL5 led to mouse infertility, and showed that TCFL5 potentially acts as a DNA/RNA-binding protein to regulate germ cell development at the transcriptional and post-transcriptional levels.

## RESULTS

### Localization of TCFL5 in mouse testes

Since there are contradictory reports of differences between the expression and localization of TCFL5 RNA and protein (Maruyama et al., 1998; Shi et al., 2013; Siep et al., 2004), we analyzed the spatiotemporal expression of TCFL5 in mouse testes and showed that TCFL5 protein was expressed exclusively in the mouse testes—with two bands at 50 kDa and 55–70 kDa (Fig. 1A). The expression of TCFL5 in mouse testes was also dramatically elevated, starting from P21 and continuing into adulthood (Fig. 1B). Consistent with this developmental patten, we also uncovered a preferential location for TCFL5 protein in pachytene spermatocytes and round spermatids—subpopulations of germ cells that we purified through the STA-PUT velocity sedimentation method we published previously(Wang et al., 2019) (Fig. 1C). Next, we immunostained for TCFL5 and determined that the TCFL5 signals were located in middle-and late-pachytene spermatocytes at stages V and VI in the seminiferous tubules of mouse testes; however, TCFL5 was absent in early-pachytene spermatocytes at stages I–IV (Fig. 1D).

**Fig. 1.**
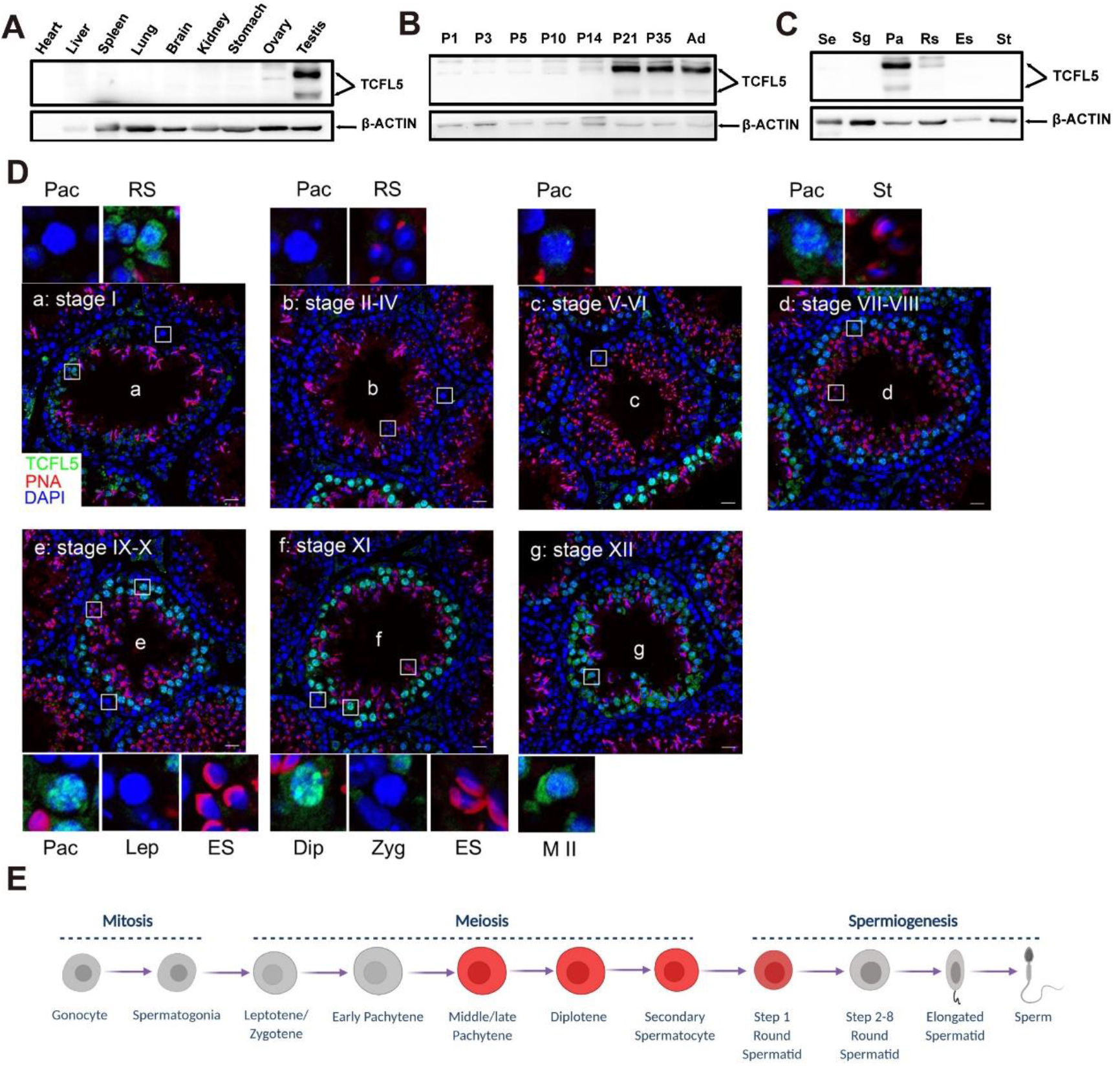
Spatiotemporal expression of TCFL5 in mouse tissue. Western blot analysis of TCFL5 protein in lysates from adult mouse tissues (A). Testicular tissues collected at different time-points during postnatal development (B), and isolated spermatogenic cell populations—including Sertoli cells (Se), spermatogonia (Sg), pachytene spermatocytes (Ps), round spermatids (Rs), elongated spermatids (Es), and spermatozoa (St) (C). β-Actin served as an internal loading control. (D) Immunostaining of testicular sections from adult wild-type mice for TCFL5 (green) and PNA (red); PNA is an acrosomal marker and DNA was stained with DAPI. Image magnifications were taken in the focused areas as shown. The spatiotemporal expression of TCFL5 in germ cells is shown by the schematic diagram (E). Pac, pachytene spermatocytes; Rs, round spermatids; St, spermatozoa; Lep, leptotene spermatocytes; Es, elongated spermatids; Dip, diplotene spermatocytes; Zyg, zygotene spermatocytes; M II, second meiosis (scale bars, 20 μm).

Notably, TCFL5 signals were present in both nucleus and cytoplasm of pachytene spermatocytes at stage VII–VIII in sections of seminiferous tubules (Fig. 1D), and the signals shifted to the nucleus in pachytene spermatocytes at stages IX and X, as well as diplotene spermatocytes at stage XI (Fig. 1D). The signals again appeared in both nuclear and cytoplasmic regions of secondary spermatocytes in meiosis II (MII) and in early round spermatids at stage I (Fig. 1D). This distinct expression pattern suggested that the actions of TCFL5 may be related to *in vivo* function during the transition from late-meiotic to post-meiotic stages in mouse spermatogenesis (as shown in Fig. 1E).

### Infertility in *Tcfl5*-heterozygous mice

To elucidate the *in vivo* function of TCFL5, we generated *Tcfl5*-knockout mice using CRISPR-Cas9 technology-mediated gene targeting (Fig. S1A). We designed small guide RNAs (gRNAs) to target exons 1–4 and deleted these four exons (and the introns between them, Fig. S1A). The first generation (F1) of *Tcfl5* mutants was successfully sired from four founder lines with either indels or deletions of *Tcfl5* alleles (Fig. S1A and S1B). Testicular sizes and body-weight ratios were identical in *Tcfl5^+/+^* and *Tcfl5^−/−^* allelic mice (Fig. 2A and Fig. S1C), and western blots and immunohistochemical staining revealed a diminution in TCFL5 protein levels in *Tcfl5*^+/−^ mouse testes in comparison to *Tcfl5*^+/+^ mice (Fig. 2B and Fig. 2C). However, we were unable to generate mice with *Tcfl5^−/−^* alleles from any F1 heterozygous fathers derived from the four founders according to the over-one-year breeding experiments, as all male *Tcfl5* mice with heterozygous alleles were likely to be infertile (Fig. 2D).

**Fig. 2.**
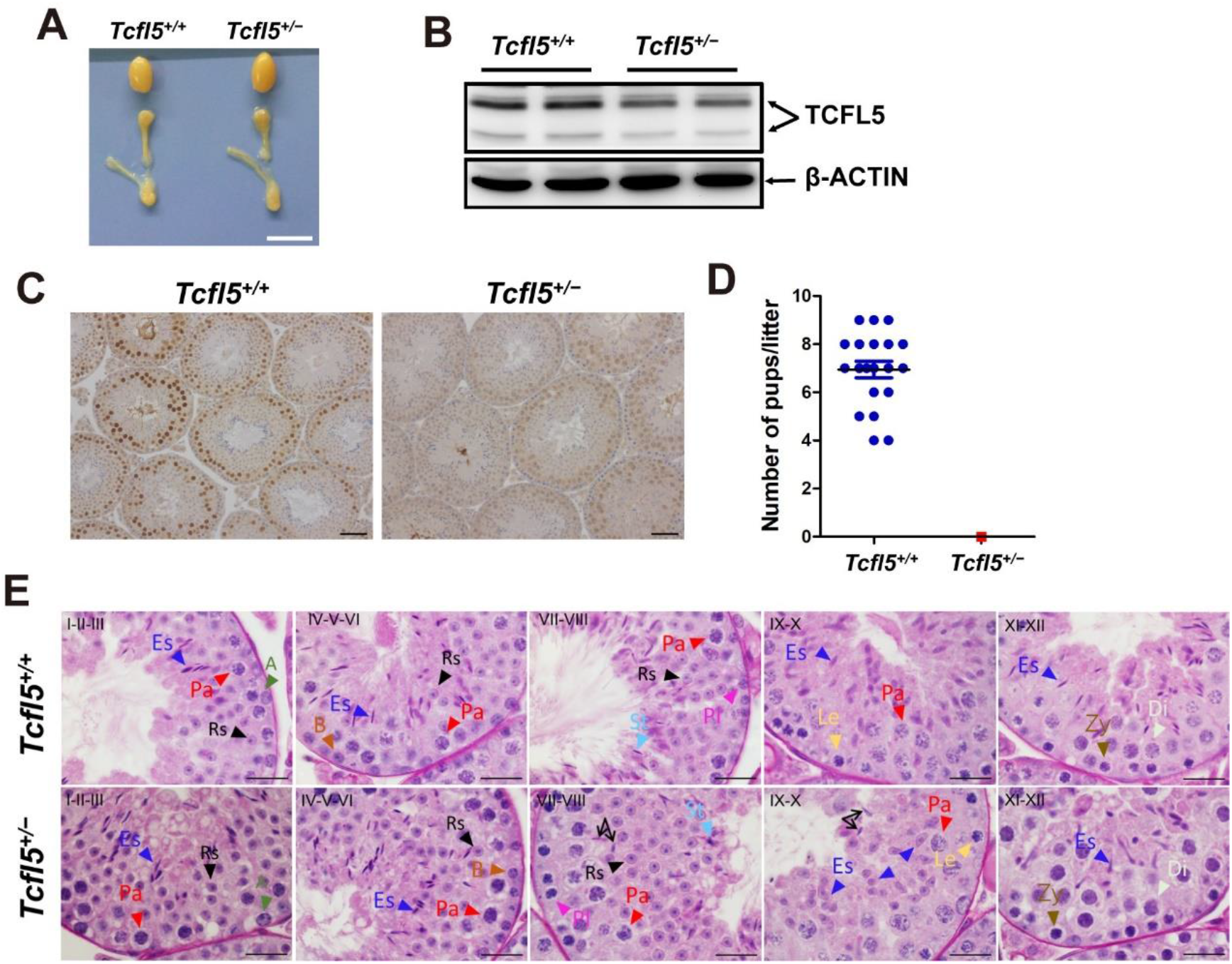
Generation of *Tcfl5* mutant mice by CRISPR/Cas9. (A) General morphology of testes and epididymides in adult *Tcfl5*^+/+^ and *Tcfl5*^+/−^ mice. (B, C) Western blot analysis and immunohistochemical staining confirmed the decrease in TCFL5 protein in adult *Tcfl5*^+/−^ mouse testes (with β-actin as the loading control). (D) Breeding experiments between *Tcfl5*^+/+^ and *Tcfl5*^+/−^ males. A mouse from each genotype was coupled with mates of normal fertility. The results were determined using nine cages of *Tcfl5*^+/−^ male mice from all four lines (see Fig. S1A) (data are presented as mean ± s.d.; dots in Fig. 2D represent 20 offspring litters derived from *Tcfl5*^+/+^ but none for *Tcfl5*^+/−^ mice; *** indicates P < 0.001 by unpaired Student’s *t test)*. (E) Morphology within the 12 stages governing the growth and development of spermatogenic epithelium. The 12 stages of seminiferous epithelium (top left) were identified according to PAS staining and cell-arrangement patterns. A, type A spermatogonia; B, type B spermatogonia; Pl, preleptotene spermatocytes; Le, leptotene spermatocytes; Zy, zygotene spermatocytes; Pa, pachytene spermatocytes; Di, diplotene spermatocytes; Rs, round spermatids; Es, elongated spermatids; St, spermatozoa (scale bars, 50 μm).

Next, our histologic analysis revealed extensive defects in testicular seminiferous tubules and epididymal tubules in *Tcfl5*^+/−^ mice compared with *Tcfl5^+/+^ mice*; i.e., the numbers of elongating spermatids were significantly decreased in *Tcfl5*^+/−^ seminiferous tubules and epididymal tubules, multinucleated cells appeared around the lumen of these seminiferous tubules, and we observed many degenerating cells with round-shaped nuclei in epididymal tubules—as indicated by the black arrowhead (Fig. S1D).

### Impaired spermiogenesis in testes of *Tcfl5*^+/−^ mice

To investigate the spermatogenic defects in the seminiferous tubules of *Tcfl5*^+/−^ mice, we conducted periodic acid-Schiff (PAS) and hematoxylin staining (Fig. 2E), and double-immunostaining with peanut agglutinin (PNA, a sperm acrosome marker) and DAPI (a nuclear indicator, Fig. 3A) to reveal changes in spermatogenesis and defects in spermiogenesis. In contradistinction to the seminiferous epithelium of *Tcfl5*^+/+^ mice in which the arrangements of spermatogonia, spermatocytes, round spermatids, and elongating spermatid layers were normal from the base to the lumen (Fig. 3A), *Tcfl5*^+/−^ mouse testes exhibited extensive defects during the 12 stages encompassing spermatogenic epithelium. For example, we observed that the deformation of round spermatids into elongating spermatids was apparently impaired at stages IX and X (blue arrowhead, Fig. 2E); that the nuclei of steps 9 and 10 elongating spermatids were irregular in size and shape; that the subsequent elongating spermatids at steps 11 and 12 presented with morphologic abnormalities (Fig. 3A); and that elongating spermatids at step 16, which would normally be released into the lumen, still remained in the epithelium of stages VII and VIII and stages IX and X (black arrows, Fig. 2E). Compared with *Tcfl5*^+/+^ mice, elongating spermatids at steps 9 and 10 of *Tcfl5*^+/−^ mice manifested malformed acrosomes. For example, the nucleus was over extended and its structural integrity (such as hooked head and dorsal angle) was lost; the acrosome did not extend normally upon losing its dorsal and ventral fins, and therefore all elongating spermatids in the following steps (11–16) were globally malformed (as illustrated in Fig. 3B); and elongating spermatids that should not be present at steps 9 and 10 were still found at stage I of the seminiferous epithelium (Fig. 3A). By contrast, all types of germ cells prior to step-9 elongating spermatids in *Tcfl5*^+/−^ mouse testes were extant and appeared normal in the epithelium (Fig. 2E and Fig. 3A).

**Fig. 3.**
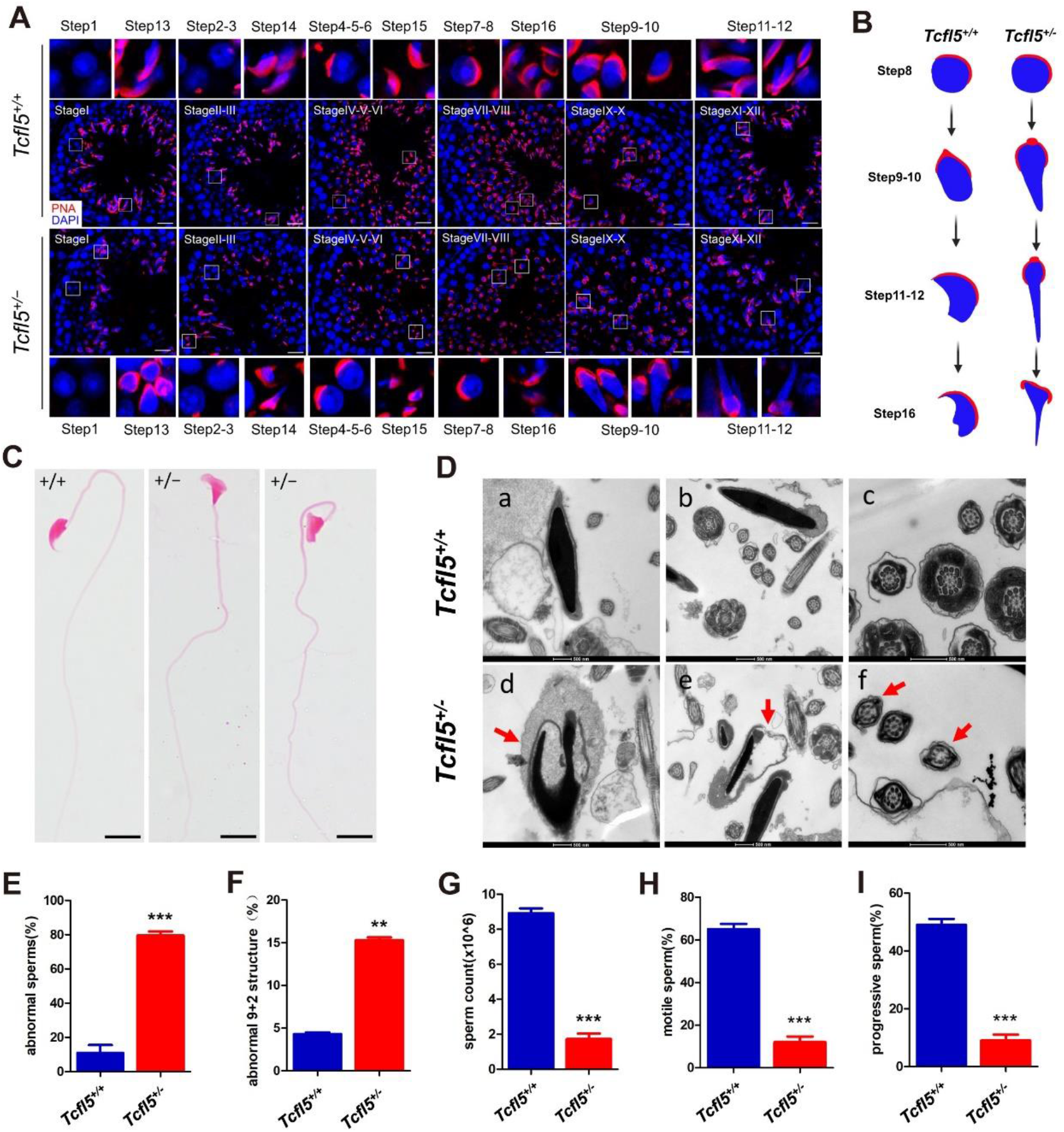
Defects in spermiogenesis between *Tcfl5*^+/+^ and *Tcfl5*^+/−^ mice. (A) PNA (red)-and DAPI (blue)-stained testis sections from adult *Tcfl5*^+/+^ and *Tcfl5*^+/−^ mice. The seminiferous tubule stage was identified based on the acrosomal morphology marked by PNA and arrangement of spermatogenic cells, and the 16-step spermiogenesis process of round and elongated spermatids was identified by the PNA-signal pattern (scale bars, 20 μm). (B) Schematic diagram showing the key steps of the differences in sperm deformation between *Tcfl5*^+/+^ and *Tcfl5*^+/−^ mice. (C) Sperm from *Tcfl5*^+/+^ and *Tcfl5*^+/−^ mouse epididymides stained by eosin showing abnormal morphologies. (D) Electron microscopic analysis of epididymal sperm from *Tcfl5*^+/+^ and *Tcfl5*^+/−^ mice, with red arrows indicating the head and tail of the sperm of the *Tcfl5*^+/−^ mice (scale bars, 500 nm). (E) Sperm with abnormal morphologies (data are presented as mean ± s.d., n=3, *** indicates p< 0.001 by unpaired Student’s *t test*) and(F) the percentages of sperm with an abnormal “9+2” structure were also determined (data are presented as mean ± s.d., n=2, ** indicates p< 0.01 by unpaired Student’s *t test*). Comparison of sperm count (G), motile sperm (H), and progressive sperm (I) in *Tcfl5*^+/+^ and *Tcfl5*^+/−^ mouse epididymides by CASA (data are presented as mean ± s.d., n=3 per group, *** indicates p value <0.001 by unpaired Student’s *t test*).

We then evaluated the sperm parameters and found that the proportion of head malformations of mature sperm in *Tcfl5*^+/−^ mouse epididymides was higher (79.5% ± 4.2) compared to the proportion in *Tcfl5*^+/+^ mice (10.9% ± 7.9) (p<0.001, unpaired t test, Fig. 3C and 3E). To further examine the defects in *Tcfl5*^+/−^ mouse sperm, we performed transmission electron microscopy (TEM) to visualize sperm ultrastructure (Fig. 3D). In contrast to the *Tcfl5*^+/+^ mouse sperm that exhibited long oval shapes with their acrosomes attached to the nuclei in the integrated “9 + 2” arrangement of microtubules within their tails (a–c in Fig. 3D), the nuclei of *Tcfl5*^+/−^ sperm were severely deformed and the acrosomes were detached from the nuclei (d, e in Fig. 3D), and some outer doublet microtubules were missing (indicated by the red arrows, f in Fig. 3D). The percentage of microtubules missing in *Tcfl5*^+/−^ mouse sperm tails was significantly higher relative to *Tcfl5*^+/+^ mice (p<0.01, unpaired *t* test, Fig. 3F). In addition, the percentages of motile and progressively motile sperm from *Tcfl5*^+/−^ mice were significantly lower than in *Tcfl5*^+/+^ mice (p<0.001, unpaired *t* test, Fig. 3H and I). Overall, we concluded that there was a gene haploinsufficiency in *Tcfl5*^+/−^ mice that prevented spermiogenesis at stages IX and X during the deformation of round spermatids to elongating spermatids.

### TCFL5 activates the expression of genes functionally related to mouse spermatogenesis

To investigate the molecular events underlying the spermatogenic defects in *Tcfl5*^+/−^ mouse sperm, we performed high-throughput RNA sequencing (RNA-seq) on purified pachytene spermatocytes in which TCFL5 protein appeared to be preferentially localized. Using adult wild-type and *Tcfl5*^+/−^ mouse testes, we collected pachytene spermatocytes purified to over 80% (Fig. S2A and S2B). The comparison of mRNA expression profiles revealed that 260 and 458 genes were significantly up-and down-regulated, respectively (Fig. 4A) in *Tcfl5*^+/−^ vs. wild-type pachytene spermatocytes, as shown in the scatterplot (Fig. 4B; p < 0.05, fold-change >2; see detailed data in Table S1 and in the GEO database, accession no. GSE176240). STRING protein network analysis of downregulated genes revealed that these transcripts were associated with cell development, cell differentiation, intercellular adhesion, and spermiogenesis (Fig. S2E). Our heatmap depicts the downregulated genes associated with spermiogenesis and spermatogenesis (Fig. 4C and Fig. S2F; p <0.05, fold-change >2). The attenuated expression of the genes associated with intercellular adhesion may partially explain the inability of sperm at stages VII–X to be released from the epithelium in a timely manner.

**Fig. 4.**
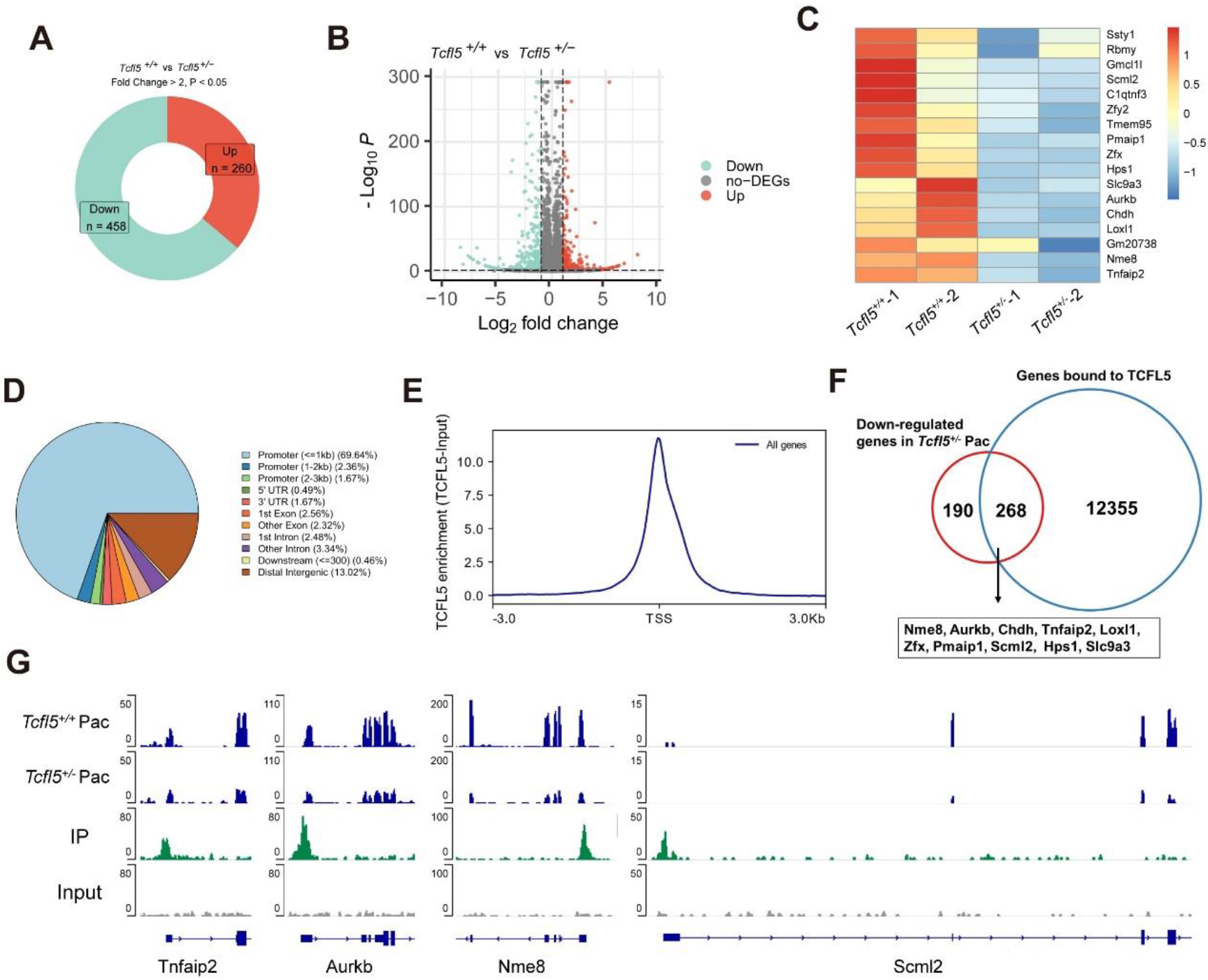
Transcriptional activation by TCFL5 of gene expression in mouse spermiogenesis. (A) Genes up-or down-regulated using high-throughput RNA sequencing between *Tcfl5*^+/+^ and *Tcfl5*^+/−^ mouse testes. Three independent samples were collected from each genotype and genes manifesting at least a two-fold change and P < 0.05 were selected for comparison. (B) Scatterplot of differentially expressed transcripts between *Tcfl5*^+/−^ and *Tcfl5*^+/+^ pachytene spermatocytes were isolated from adult *Tcfl5*^+/+^ and *Tcfl5*^+/−^ mice with the STA-PUT method (*Tcfl5*^+/+^ n= 2; *Tcfl5*^+/−^ n= 2). Each dot represents a gene that was significantly changed (fold-change >2, P <0.05). (C) Heatmap of genes that are differentially up-and down-regulated by TCFL5 and that are closely related to spermatogenesis. (D) Distribution map of the proportions of potential binding positions of genes found in the TCFL5 ChIP experiments. (E) Prediction of the distribution area of TCFL5-binding sites at or near the targeting gene transcription start sites (TSS); TCFL5 was enriched at the regions from 3 kb to 3 kb relative to the TSS. (F) ChIP and RNA-Seq experiments suggested that 268 genes (58.5%) of the 458 down-regulated genes were directly regulated by the transcriptional regulation of TCFL5. (G) Genome browser view of TCFL5 ChIP-seq and RNA-seq reads on Tnfaip2, Aurkb, Nme8, and Scml2 gene loci.

To identify genes directly regulated by TCFL5, we next performed chromatin immunoprecipitation with massively parallel DNA sequencing (ChIP-seq) to identify the binding of TCFL5 by using P-21 testes (in which TCFL5 was abundantly expressed). The enrichment of TCFL5 protein was verified by western immunoblotting (Fig. S4C). Model-based analysis of ChIP-seq(Zang et al., 2009) identified 12,623 peak-associated genes with significant enrichment by TCFL5 (Table S2). We then analyzed the ChIP-seq data by mapping DNA sequences to genomic regions and found that TCFL5 bound to the promotor regions of genes as expected (Fig.4D), and that a majority of TCFL5-binding sites were close to the center of the transcription start sites (TSS, Fig. 4E). The distribution of TCFL5-precipitated reads on gene bodies thus indicated that TCFL5 is a potential transcriptional activator in mouse testes (Fig. S2D).

We analyzed RNA-seq data in combination with the ChIP-seq data and identified 268 downregulated genes, including spermatogenesis-associated genes (as listed in Fig. 4F). A genome-browser view revealed that TCFL5 bound to the promoter regions of *Tnfaip2*, *Aurkb*, *Nme8,* and *Scml2*, and that it activated their expression (Fig. 4G). We noted that *Scml2* was one of the genes that showed a pronounced reduction in transcription (Table S1, Fig. 4F; and q-PCR data in Fig. S2F). To evaluate the regulation of *Scml2* by TCFL5, we constructed a Tcfl5*-*expression vector by inserting Tcfl5 CDS into a pcDNA3.1+ vector, and determined the peak sequence in which TCFL5 bound to the *Scml2* promoter region using a dual-luciferase assay by co-transfecting 293T cells with a Tcfl5 expression plasmid, a pGL3 basic plasmid-containing peak sequence, and a *Renilla* luciferase vector. Our results established that relative luciferase activity increased significantly in the experimental group compared to the control groups (Fig. S2G). In addition, western blotting analysis indicated an attenuated expression of SCML2 protein in *Tcfl5*^+/−^ mouse testes (Fig. S2H). These results indicated that TCFL5 activated the expression of *Scml2* and other genes that are related to spermatogenesis by binding to their promoters (Fig. S3B).

### TCFL5 interacts with the RNA-binding proteins FXR1 and DHX9

To further scrutinize the regulatory mechanism(s) subserving TCFL5 function in spermatogenesis, we profiled TCFL5-interacting proteins with TCFL5 antibody immunoprecipitation (IP) using lysates of mouse testes on P21. In two independent IP-mass-spectrometric experiments, we identified 57 candidate proteins that may interact with TCFL5 (Fig. S3A and Table S3). Intriguingly, we found that a majority of the interacting proteins of TCFL5 were functionally related to RNA and posttranscriptional processing—including heterogeneous nuclear ribonucleoproteins (HnRNPs), splicing factor-related proteins, ribosomal proteins, and some classical, well-studied RNA-binding proteins (i.e., FXR1 and DHX9)—as shown by Coomassie blue-staining prior to sequencing by mass spectrometry (Fig. 5A). Gene-ontology (GO) analysis displayed terms significantly related to mRNA processing, RNA splicing, and/or mRNA stability (Fig. 5B). These results suggested that different regions of TCFL5 may be also responsible for binding to DNA and RNA with or without other proteins, and that TCFL5 may possess dual DNA-and RNA-binding capacities. If TCFL5 binding to DNA were stabilized by its binding to RNA, then RNase treatment of chromatin should reduce TCFL5 occupancy. To test this idea, we treated nuclear chromatin with RNase inhibitor or RNase A treatment, and, indeed, the levels of TCFL5 bound to chromatin were significantly diminished when the chromatin preparation was treated with RNase A—consistent with the concept that RNA may contribute to the stability of TCFL5 in chromatin (Fig. 5C).

**Fig. 5.**
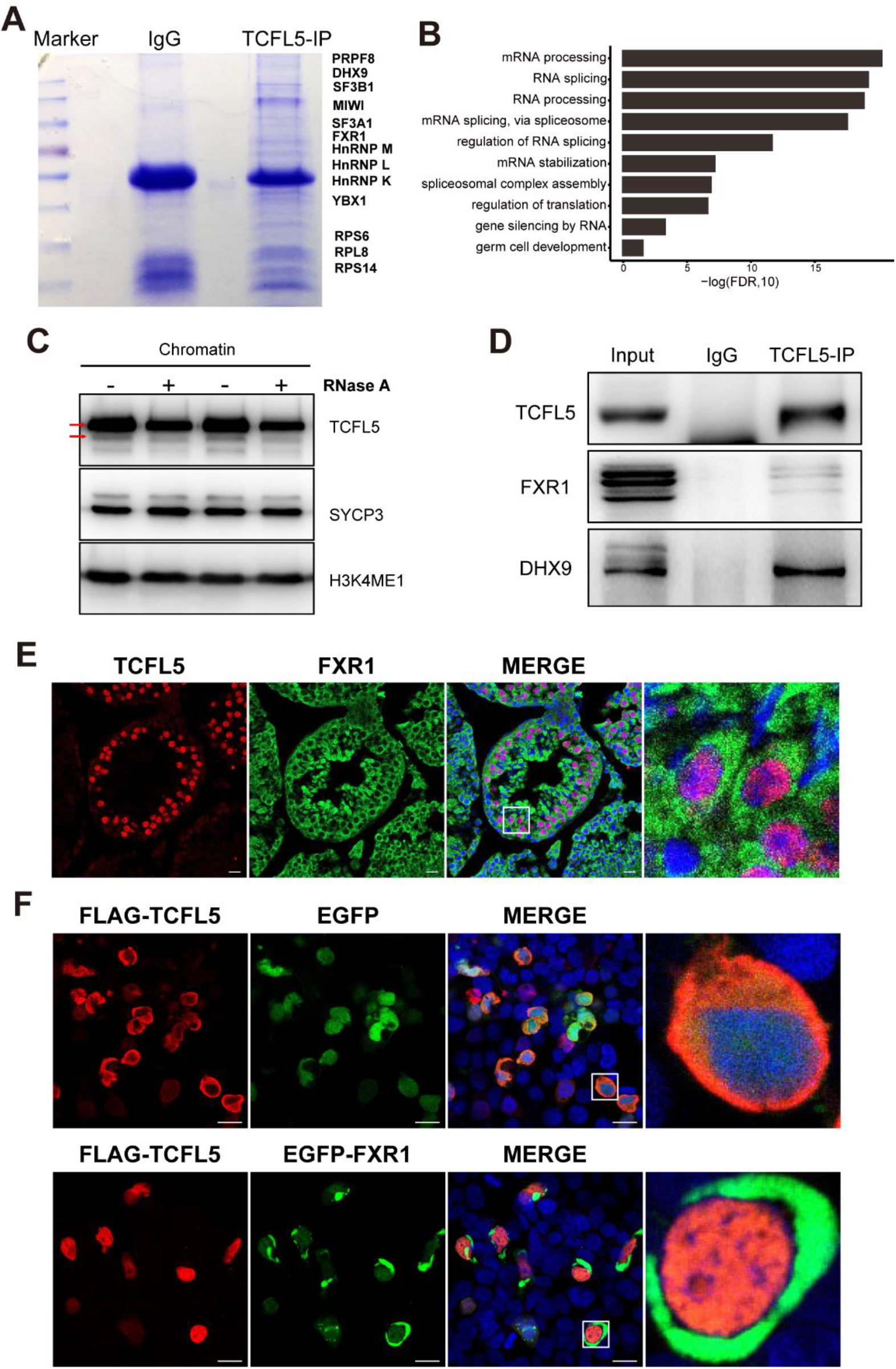
Interaction of TCFL5 with FXR1 and DHX9. (A) Coomassie blue staining after TCFL5 immunoprecipitation in 21-d mouse testes indicating the differentially enriched protein positions as revealed by protein mass spectrometry. (B) Gene ontology (GO) term-enrichment analysis of 57 candidates (see Fig. S3) obtained from two independent TCFL5 immunoprecipitation and mass-spectrometry experiments. (C) Chromatin-binding assay as assessed by western blotting analysis showing the levels of TCFL5, SYCP3, and H3KEME1 in a nuclear chromatin preparation after RNase A inhibitor or RNase A treatment. (D) Immunoprecipitation followed by western blotting in 21-d mouse testes confirming the interaction of TCFL5 with FXR1 and DHX9. (E) Co-localization of TCFL5 and FXR1 in the seminiferous tubules of mouse testes. (H) Comparison of the co-expression of TCFL5-FLAG protein and EGFP protein (upper panel) and TCFL5-FLAG protein and FXR1-EGFP protein (lower panel) in 293T cells after in vitro transfection

Of 57 candidate proteins that interacted with TCFL5, we further confirmed at least two known dual DNA-and RNA-binding proteins—FXR1 (fragile-X mental retardation syndrome-related protein 1) and DHX9 (DExH-box helicase 9)—by western blotting after TCFL5 immunoprecipitation, indicating that the interactions between/among FXR1, DHX9, and TCFL5 may coordinate transcriptional regulation in mouse testes. While there are only commercial antibodies to FXR1 available to verify its co-localization with TCFL5 in testes, we observed preferential intracellular sub-localization of FXR1 in the cytoplasm (Fig. 5E). We examined the co-expression of TCFL5-FLAG and FXR1-EGFP proteins (Fig. 5F, lower panel) in 293T cells to compare with TCFL5-FLAG and EGFP proteins (Fig. 5F, upper panel), and showed that FXR1 likely supported the transport of TCFL5 into the nucleus (Fig. 5F). Collectively, the aforementioned data indicate that TCFL5 may not only be involved in transcriptional regulation but also in RNA processing during spermatogenesis.

## DISCUSSION

Due to the scarcity of available clinical cases and detailed genetic background data— and because clinically characterizing information regarding male infertility is often lacking—animal models have been applied to examine the roles of genes and to clarify their underlying mechanism(s) of action in the processes of germ cell production (Matzuk and Lamb, 2008; Nalam and Matzuk, 2010). In the present study, we identified a new testis-specific gene, Tcfl5, that belongs to the basic helix-loop-lelix (bHLH) family, and that is required for spermatogenesis in mouse testes and for male fertility. TCFL5 is localized in middle and late spermatocytes starting from stage V to the round-spermatid stage in neonates and demonstrates both nuclear and cytoplasmic expression. We observed the signal intensity of TCFL5 to be weak in stages V–VIII spermatocytes, but augmented from stage IX onward, with signal intensity in the nucleus stronger than in the cytoplasm. By contrast, intensity in these compartments was reversed in secondary spermatocytes and round spermatids. These distinct expression patterns support the concept that TCFL5 functions as both a DNA-and RNA-binding protein from late-meiotic to post-meiotic stages.

From our RNA-seq and ChIP-seq results, it was obvious that TCFL5 bound to at least 258 genes (58.5% of the downregulated genes by RNA-Seq), validating TCFL5’s transcriptional regulation of genes associated with germ cell development and serving as a critical factor in the mouse testis. Scml2 in the testis has been reported to deubiquitinate RNF8-dependent monoubiquitination of histone H2A at lysine 119 (H2AK119ub), thus inhibiting the establishment of H3K27 acetylation(Adams et al., 2018; Hasegawa et al., 2015). H3K27 acetylation is a marker of active enhancers and is followed by H3K4 dimethylation—a modification associated with active transcription(Creyghton et al., 2010; Talasz et al., 2005). We showed that TCFL5 bound to the promoter region of *Scml2* and regulated the transcription of *Scml2*. Therefore, the decrease in *Scml2* expression in *Tcfl5* mutants may have affected the reactivation of male reproductive genes after meiosis, and ultimately led to the production of abnormal sperm. Tnfaip2 and Nme8 are components of the acrosome and fibrous sheath, respectively(Miranda-Vizuete et al., 2003; Smith et al., 2013; Wolf et al., 1994). *Nme8* mutation is associated with accelerated loss of sperm motility and impaired protamination of sperm chromatin(Smith et al., 2013). *Aurkb* mutant mice exhibited reduced sperm counts, and in some cases complete absence of spermatozoa, manifesting multinucleated cells in the seminiferous tubules and subfertility defects(Kimmins et al., 2007). AURKB is a chromosomal passenger protein That regulates kinetochore-microtubular attachments and cytokinesis, preventing tetraploidization(Adams et al., 2001; Steigemann et al., 2009), and this may explain the phenotype of multinucleated cells present in *Tcfl5*^+/−^ mouse testes. TCFL5 may also regulate spermatogenesis by activating Tnfaip2, Nme8, and Aurkb expression.

Proteins that bind DNA or RNA are often considered and studied independently of one another. However, recent studies have indicated that about 8% of nuclear proteins possess dual DNA-and RNA-binding capacities, and many transcriptional factors are capable of binding diverse types of RNA(Hudson and Ortlund, 2014). For example, although the DNA-binding protein CTCF was initially described as a conserved transcription factor(Filippova et al., 1996), it was subsequently also reported to be a specific RNA-binding protein with a previously unrecognized RNA-binding region(Kung et al., 2015; Saldaña-Meyer et al., 2014) and shown to be involved in alternative spicing(Shukla et al., 2011). The classical transcription factor Yy1 also exhibits RNA-binding function(Jeon and Lee, 2011; Sigova et al., 2015). The binding of yy1 to RNA transcripts associated with regulatory elements can enhance its occupation at these elements(Jeon and Lee, 2011; Sigova et al., 2015), and the chromatin-associated protein PARP1 associates with nucleosomes that harbor exonic H3K4me3 marks, binds nascent RNA, and regulates splicing co-transcriptionally by recruiting splicing factors(Matveeva et al., 2016; Melikishvili et al., 2017). Transcription factor Sox2 can associate with RNA while binding to its cognate DNA sequence to form ternary RNA/Sox2/DNA complexes that co-transcriptionally regulate RNA metabolism during somatic cell reprogramming(Hou et al., 2020). In addition, some RNA-binding proteins localize to active chromatin regions and directly participate in transcriptional control, with active gene promoters constituting the major binding regions(Xiao et al., 2019). SR proteins comprise a family of RNA-binding proteins involved in both constitutive and regulated splicing(Lin and Fu, 2007), and SR protein SRSF2 was reported to be a transcriptional regulator associating with gene promoters as part of the 7SK complex and activating transcription via promoter-proximal nascent RNA(Ji et al., 2013). We discovered that TCFL5 interacts with many RNA-binding proteins according to our mass-spectrometric results, and therefore speculate that TCFL5 exerts RNA-binding actions and is involved in post-transcriptional regulation.

Our chromatin-binding experiment also showed that after the removal of RNA by RNase A, the chromatin binding by TCFL5 protein was reduced—indicating that TCFL5 binds DNA in an RNA-dependent manner, and also confirms the overall RNA-binding function of TCFL5. Of the plethora of interacting proteins, the first group was composed of hnRNPs, i.e., HnRNP M, HnRNP L, and HnRNP K. HnRNPs represent a large family of RNA-binding proteins that contribute to multiple aspects of nucleic acid metabolism—including alternative splicing, mRNA stabilization, and transcriptional and translational regulation(Geuens et al., 2016). The second group comprised splicing factors, i.e., SF3B1, SF3A1, and PRPF8, while the third consisted of ribosomal proteins (e.g·., RPS6, RPL8, and RPS14). In addition, there were also some classical RNA-binding proteins such as DHX9 and FXR1. DHX9 is a member of the DExD/H-box family of helicases and was initially portrayed as a transcriptional co-activator acting as a bridging factor between the transcriptional co-factor CREB-binding protein (CBP)/p300 and Pol-II, recruiting the latter to the CREB/CBP/p300 complexes at the promoter(Nakajima et al., 1997). FXR1 was also reported to regulate transcription by recruiting transcription factor STAT1 or STAT3 to gene promoters at the chromatin interface, and at least partially mediating cellular proliferation(Fan et al., 2017). Thus, TCFL5 co-participates in transcriptional activation of target genes through interactions with DHX9 and FXR1 (Fig. S3B). However, in addition to transcriptional regulation, DHX9 is necessary for post-transcriptional regulation—including participating in the translation of viral and JUND mRNA(Hartman et al., 2006; Ranji et al., 2011), pre-mRNA splicing(Hartmuth et al., 2002; Zhang et al., 1999), and RNA transport by binding directly to F-actin and HnRNP C1(Zhang et al., 2002). FXR1 modulates *p21/Cdkn1a/Cip1/Waf1* mRNA stability in myoblast cell-cycle progression(Davidovic et al., 2013), operates in dendritic mRNA transport(Dictenberg et al., 2008), and activates translation(Vasudevan and Steitz, 2007; Vasudevan et al., 2007). We therefore speculated that TCFL5 is involved in RNA splicing, mRNA stability regulation, and transport and translational regulation through its interactions with DHX9 and FXR1 (Fig. S3B), and thus TCFL5 exerts crucial actions in the overall mechanisms involved in spermatogenesis.

Collectively, we postulate that TCFL5 plays an important role in spermiogenesis via two pathways—one in the regulation of *Scml2*, *Tnfaip2*, *Aurkb,* and *Nme8* gene expression, and the other in post-transcriptional regulation by interacting with RNA-binding proteins such as DHX9 and FXR1 (Fig. S3B). Because of the important functions ascribed to TCLF5, we posit that *Tcfl5* haploinsufficiency leads to dysfunctions in spermatogenesis. These data thus provide novel clues toward understanding spermatogenesis and in the treatment of male infertility.

## MATERIALS AND METHODS

### Generation of Tcfl5-knockout mice by CRISPR/Cas9

Exons 1–4 of the *Tcfl5* gene were targeted with four sgRNAs, and the synthesized paired oligonucleotides for sgRNAs were annealed and cloned into the pUC57-sgRNA expression vector (51132, Addgene). Oligo sequences were as follows: sgRNA1 up, taggTGGCCGGCGCGTCGTCTG; sgRNA1 down, aaacCAGACGACGCGCCGGCCA; sgRNA2 up, taggCGGGCGCGACGGGCGGCG; sgRNA2 down, aaacCGCCGCCCGTCGCGCCCG; sgRNA3 up, taggCCTTCAGGTTAGTTGGCA; sgRNA3 down, aaacTGCCAACTAACCTGAAGG; sgRNA4 up, taggTAGCTTCGTATTGTTTTG; and sgRNA4 down, aaacCAAAACAATACGAAGCTA. The Cas9 expression plasmid (44758, Addgene) was linearized with *Age* I and used as the template for in vitro transcription using the T7 Ultra Kit (AM1345, Ambion). sgRNA expression plasmids were linearized with *Dra* I and used as templates for in vitro transcription using the MEGAshortscript Kit (AM1345, Ambion). Transcribed Cas9 mRNA and sgRNA were both purified by using the MEGAclear Kit (AM1908, Ambion). C57BL/6 mice were purchased from Beijing Vital River Laboratories animal center and housed in standard cages and maintained on a 12-h light/dark cycle with food and water ad libitum. The microinjection of fertilized mouse eggs was performed in the Zhang Laboratory, Institute of Laboratory Animal Science, the Chinese Academy of Medical Sciences. In brief, fertilized eggs were injected with a mixture of Cas9 mRNA (12 ng/μl) and four sgRNAs (6 ng/μl each). Founder mice were backcrossed to C57BL/6J. The PCR primers for genotyping were as follows: forward primer (P1), GCTTCATCAGACGCTTGGTC; reverse primer (P2), TGCATGATGTGTGGAGCTGACAGC; forward primer (P3), CAGTGGGCGTGTCTTTATGAG; and reverse primer (P4), CTTGCTACTTAAGACTGGC. Primers P1 and P2 were used for wild-type and primers P3 and P4 for the mutant. The wild-type allele generated a 407-bp band while the knockout allele produced a 5160-bp band. Mice were maintained under specific pathogen-free (SPF) conditions according to the guidelines of the Institutional Animal Care and Use Committee of Nanjing Medical University.

### Isolation of pachytene spermatocytes

We isolated pachytene spermatocytes from adult mice using the STA-PUT method. Mice testes were first digested with collagenase IV (1 mg/ml), and the dispersed seminiferous tubules were washed with DMEM and then centrifuged at 500 *g*. The pellet was further digested with 0.25% trypsin-containing DNase I (1 mg/ml), and we terminated digestion with 10% FBS. The resulting cellular suspension was filtered with a 40-μm filter to prepare a single-cell suspension, which was then loaded onto a cell-separation apparatus (ProScience, Canada) using a 2–4% bovine serum albumin (BSA) gradient (2% and 4% BSA were loaded onto the separation-apparatus chamber). After 2–3 h of sedimentation, we harvested the cell fractions and identified pachytene spermatocytes according to their morphologic characteristics, cell diameters, and Hoechst staining under light microscopy (Ti2-U, Nikon, Japan).

### Western immunoblotting analysis

Testes and cultured 293T cells were lysed in RIPA buffer (50 mM Tris-HCl, 150 mM NaCl, 1% NP40, 1 mM DTT, 0.5% sodium deoxycholate, 0.05% SDS, and 1 mM EDTA) containing protease-inhibitor cocktail (11697498001, Roche). Protein lysates were separated on 10% SDS-PAGE gels followed by electrotransfer onto PVDF membranes. The blots were blocked for 2 h with 5% nonfat milk and then incubated overnight at 4°C with the following primary antibodies: anti-TCFL5 (1:1000; ab188075, Abcam), anti-SCML2 (1: 1000; kindly provided by Dr. Mengcheng Luo, Wuhan University, China), anti-FLAG a (1:1000; F1804, Sigma), anti-FXR1 (1:1000; A8697, Abclonal), and anti-DHX9 (1:1000; A17955, Abclonal). HRP-conjugated secondary antibody (1:2000; ZB2301, Zhongshan Jinqiao Biotechnology) and ECL reagent were used to visualize immunopositive bands.

### Immunofluorescence

Testes were fixed in 4% paraformaldehyde at 4°C, dehydrated in sucrose, embedded in OCT, and then cut at a 5-μm thickness using a cryostat microtome (Cryotome FSE, Thermo Fisher Scientific). We executed immunostaining on testicular cryosections with primary antibody (anti-TCFL5), and Cy™2-conjugated secondary antibody (1:200; 711-225-152, Jackson ImmunoResearch, USA) was used for visualization of immunostaining. For acrosomal staining, cryosections were incubated for 1 h at 37°C with rhodamine-conjugated PNA (1:2000; RL-1072, Vector Laboratories), and nuclear DNA was stained with DAPI (1:500; D9542, Sigma). Stained testis sections were visualized on a confocal microscope (LSM800, Carl Zeiss).

### Immunocytochemistry

The Fxr1 CDS was cloned into a pEGFP-C1 vector and co-expressed with EGFP TAG (the co-expression plasmid was constructed by Tsingke Biological Technology, Nanjing, China). The Tcfl5/pcDNA3.1+ and Fxr1/pEGFP-C1 plasmids were transfected into HEK 293T cells simultaneously using Calcium phosphate/CaCl2 transfection. After transfection for 36 h, the cells were fixed with 4% paraformaldehyde, permeabilized with 0.5% TritonX-100, and blocked with 2% BSA/10% normal donkey serum/0.2% TritonX-100 in PBS for 2 h at room temperature. The cells were then incubated overnight at 4°C with FLAG antibody (1:500, F1804, Sigma), and incubated with TRITC-conjugated secondary antibody (1:200, 715-025-150, Jackson ImmunoResearch, USA) for 2 h at room temperature. Images were taken under confocal microscopy (LSM800, Carl Zeiss).

### Histology and transmission electron microscopic analysis

Testes and epididymides were fixed in Hartman’s fixative (H0290, Sigma) for 24 h. The tissues were dehydrated with increasing concentrations of ethanol (70%, 80%, 90%, 100%), cleared in xylene, embedded in paraffin, and sectioned at 5 μm. The sections were deparaffinized, rehydrated, stained with hematoxylin and eosin (H&E; Sigma-Aldrich), or stained with PAS reagent. For transmission electron microscopic analysis, cauda epididymal sperm obtained from *Tcfl5*^+/+^ and *Tcfl5*^+/−^ mice were fixed in 2.5% glutaraldehyde in 0.2 M cacodylate buffer at 4°C overnight. Sperm were then washed with 0.2 M cacodylate buffer, dehydrated in a graded series of ethanol, and embedded and polymerized in an automated microwave tissue processor (EMAMW, Leica). Ultrathin sections were stained with uranyl acetate and lead citrate and were examined by TEM (JEM-1010, JEOL).

### Sperm analysis

Sperm from *Tcfl5*^+/+^ and *Tcfl5*^+/−^ mice were extracted and incubated in Hanks’ Balanced Salt Solution (HBSS) medium (14175-079, Gibco) at 37°C for 20 min, and samples were diluted and analyzed using Hamilton Thorne’s Ceros II system.

### Co-immunoprecipitation

For IP, whole testes from mice at postnatal day 21(P21) were lysed in RIPA buffer containing protease-inhibitor cocktail (11697498001, Roche). Protein lysates were then blocked by incubation with protein A Dynabeads (10002D, Invitrogen) for 2 h at 4°C followed by incubation with primary antibody to TCFL5 (1:50; 188075, Abcam) or control IgG (normal rabbit IgG, A7016, Beyotime) overnight at 4°C. The lysates were then incubated with protein A Dynabeads for 2 h at 4°C and proteins eluted in lysis buffer for 10 min at 100°C after stringent washing conditions in lysis buffer. The co-immunoprecipitated protein complexes were detected using western blotting with the following antibodies: anti-TCFL5 (1:1000, ab188075, Abcam), anti-FXR1 (1:1000, A8697, Abclonal), and anti-DHX9 (1:1000, A17955, Abclonal).

### Coomassie blue staining

SDS-PAGE gels were dipped in R250 dye solution for 3 h at room temperature with gentle rocking and then washed in washing buffer until the gels became transparent.

### MS spectrometry

The TCFL5 protein complexes pulled down with an anti-TCFL5 antibody from adult mice testes were analyzed by the Mass Spectrometry Core Center, Nanjing Medical University, using the entire elutions.

### RNA isolation and sequencing

Total RNA was extracted from pachytene cells taken from wild-type and *Tcfl5*^+/−^ mice using Trizol reagent (15596018, Life technologies). Two biologic replicates were executed and used for sequencing. RNAs were subjected to oligo dT selection and adaptor ligation. We performed sequencing on the BGISEQ-500 platform at BGI Tech (Shenzhen, China). Low-quality reads were filtered using the internal software SOAPnuke. Afterward, clean reads were obtained and stored in FASTQ format. The clean reads were mapped to the mouse genome (GRCm38.p5) using HISAT2. Bowtie 2 was applied to align the clean reads to the reference coding gene set, and the expression level of each gene was calculated by RSEM. Differentially expressed genes between samples were generated by DESeq2 algorithms. Fold change of ≤−2 or ≥ and adjusted q-value <0.05 were considered to show significantly differential expression.

### ChIP-seq and data analysis

Testis tissues were collected from wild-type and *Tcfl5* mutants on P21 and cross-linked with 1% formaldehyde, followed by quenching with glycine solution. Chromatin fragmentation was performed by sonication in ChIP SDS lysis buffer (50 mM Tris-HCl, pH 8; 10 mM EDTA, pH 8; and 1% SDS) using a Covaris-S220 sonicator. Cross-linked chromatin was incubated with anti-Tcfl5 antibodies in ChIP dilution buffer (16.7 mM Tris-HCL, 167 mM NaCl, 1.2 mM EDTA, 1.1% Triton X-100, and 0.01% SDS) with protease inhibitors overnight. Cross-linking was then reversed overnight at 65°C, and DNA extracted using chloroform. Purified DNA was sequenced on the BGISEQ-500 platform. Raw reads were filtered first to remove low-quality or adaptor sequences by SOAPnuke, and cleaned reads were mapped to the mouse genome (GRCm38.p5) using SOAPaligner/SOAP2. Quality control for ChIP-seq data was done with FastQC software, and peak calling for TCFL5 was carried out using Model-based Analysis for ChIP-Seq (MACS2). Genome-wide signal coverage tracks were visualized in the Integrative Genome Browser (IGV).

### Dual-luciferase reporter assay

Peak sequences from TCFL5 targets were amplified by PCR, and the Tcfl5 CDS was cloned into a pcDNA3.1+ vector and co-expressed with FLAG TAG (the co-expression plasmid was constructed by Tsingke Biological Technology, Nanjing, China). The peak sequences of target mRNAs were cloned into a pGL3 basic vector (PM0874, BioNice Co., LTD, Ningbo) using the ClonExpress MultiS One Step Cloning Kit (C113-02, Vazyme Biotech). Primers synthesized for peak sequences were as follows: Tcfl5-peak sequence, 5’-tctgcgatctaagtaagcttGTTATATGGTGGTGCTATCTTTGCTGG-3’ (forward 1) and 5’-GTTTAATCACACCCTCATAAAGACACGC-3’ (reverse 1), 5’-CTTTATGAGGGTGTGATTAAACTGGGCGTG-3’ (forward 2) and 5’-CTGCGTGTACTCCACTTCCGTCATCTC-3’ (reverse 2), 5’-CGGAAGTGGAGTACACGCAGC-3’ (forward 3) and 5’-agtaccggaatgccaagcttGACTGCCTGCCCCTTTACACTG-3’ (reverse 3); Scml2-peak sequence, 5’-tctgcgatctaagtaagcttAAAACTGCCAAAGGCTTTTTAGCATC-3’ (forward) and 5’-agtaccggaatgccaagcttCAGAGAACACTAATTTAGACACAGCAGC-3’ (reverse). PCR conditions were as follows: 95°C for 5 min; 95°C for 45 s, 60°C for 30 s, and 72°C for 1 min for 35 cycles; 72°C for 10 min, and then PCR was paused. The peak/pGL3 basic plasmid, Tcfl5/pcDNA3.1+ plasmid, and *Renilla* plasmid (graciously provided by Dr. Yunjun Xu, Nanjing Medical University, China) were transfected into HEK 293T cells simultaneously using CaCl2 transfection. After transfection for 36 h, the cells were collected and luciferase activities were determined according to the manufacturer’s instructions for the Dual-Luciferase Reporter Assay System (E1910, Promega).

### Chromatin-binding assay

The testicular tissues from two groups (100 mg/group) were prepared using 1 ml of hypotonic buffer each. One group was treated with a 1:100 dilution of RNase A (ST576, Beyotime) for 10 min at 37°C and the other was treated with 1 U/ul RNase inhibitor (N2518, Promega). For chromatin extraction, the tissue was homogenized in 1 ml of buffer A (250 mM sucrose, 10 mM Tris HCl [pH 8.0], 10 mM MgCl_2_, 1 mM EGTA, and 1X protease-inhibitor cocktail [11697498001, Roche]), and incubated for 30 min at 4°C with gentle rocking. The lysates were then passed through a 40-um cell strainer followed by centrifugation at 1200 g for 5 min at 4°C to separate nuclei (pellet) from the cytoplasmic extract (supernatant). The pellet was washed in buffer A three times and then homogenized in 400 ul of buffer B (250 mM sucrose, 10 mM Tris HCl [pH 8.0], 10 mM MgCl2, 1 mM EGTA, 1X protease-inhibitor cocktail [11697498001, Roche], 0.1% Triton X-100, and 0.25% NP-40) for 10 min at 4°C. The sample was then centrifuged at 100 g for 5 min at 4°C to remove nuclear membrane particles (pellet). The supernatant was collected and centrifuged at 1700 g for 5 min at 4°C to separate insoluble chromatin (pellet) from soluble nuclear protein (supernatant). The pellet was then washed in buffer B three times. We used 50 ul of 0.2 M HCl to release the bound protein from chromatin, and 10 ul of 1 M Tris HCl (pH 8.0) was used to neutralize the pH. The chromatin fraction was then analyzed using western blotting analysis to determine the expression of TCFL5.

### Data processing and statistical analysis

We performed a sperm CASA experiment with three experimental replicates and captured fluorescence images using confocal microscopy (LSM800, Carl Zeiss). All image data were assembled according to Adobe Illustrator (2021 edition, USA) software. Data are reported as mean ± SD unless otherwise noted in the figure legends. Significance was assigned using unpaired Student’s *t test* (* P < 0.05; ** P < 0.01; *** P < 0.001) with Excel 2016 (Microsoft, USA). We carried out the quantitative data presentation using GraphPad Prism 8.0 software (USA).

## Acknowledgments

The work was supported by the National Key Research and Development Program of China (2018YFC1003302 to X.W) and the National Natural Science Foundation of China (32070831 and 31872844 to X.W)

## Author contributions

Conception and design, W.X and X.W; collection and/or assembly of data, W. X, M.W, Y.Z, D.Q, Y.G, S.W, G.D, F.Y, L.L and S.Y; data analysis and interpretation, W.X and X.W; manuscript writing, W.X, L.L, and X.W; final approval of the manuscript, W.X and S.W; administrative support, X.W; and financial support, X.W

## Data availability statement

The data that support the findings of this study are available from the corresponding author upon reasonable request. Sequencing data have been deposited into the GEO database, accession no. GSE176240.

---

**Fig. S1.**
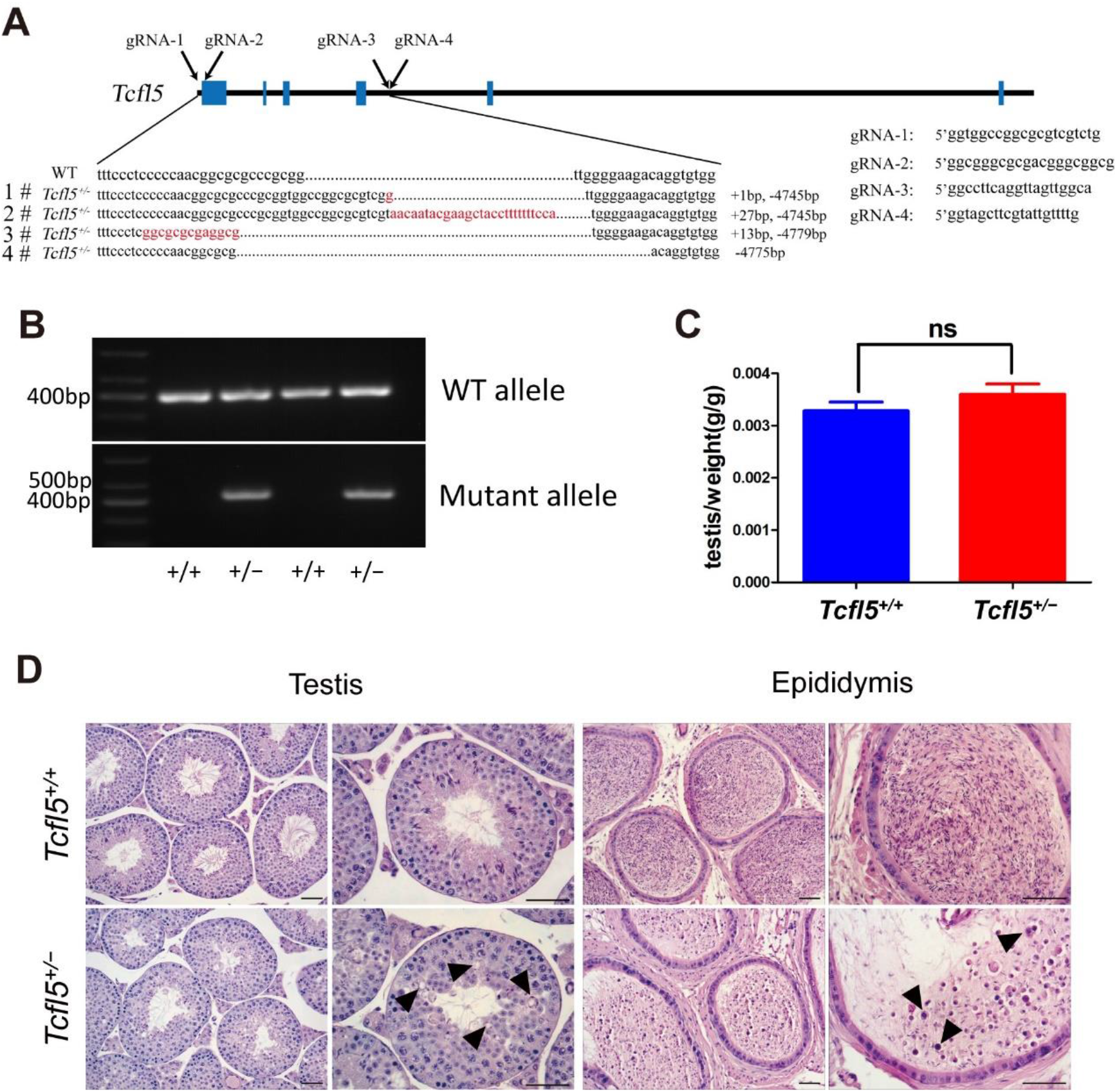
Genotyping for the presence of mutant alleles. (A) Diagram illustrating the CRISPR/Cas9 targeting strategy used for the *Tcfl5* gene in mice, including position and sequence of guide RNAs (E, exon), and (B) the presence of a band near 400 bp (lower panel) was considered to be heterozygous. (C)Testis/body-weight ratio was compared between *Tcfl5*^+/+^ and *Tcfl5*^+/−^ mice (data are presented as mean ± s.d., n=3 per group, *ns* indicates no statistical significance by unpaired Student’s *t test*). (D) Hematoxylin and eosin staining of seminiferous tubules and epididymides in adult *Tcfl5*^+/+^ and *Tcfl5*^+/−^ mouse testes. The number of elongating spermatids in the tubules of *Tcfl5*^+/−^ mice was significantly reduced, fewer mature sperm were found, and degenerating cells were observed with round-shaped nuclei. Multi-nucleated cells are depicted (black arrowheads) in *Tcfl5*^+/−^ mouse testes and cauda epididymides (scale bars, 50 μm).

**Fig. S2.**
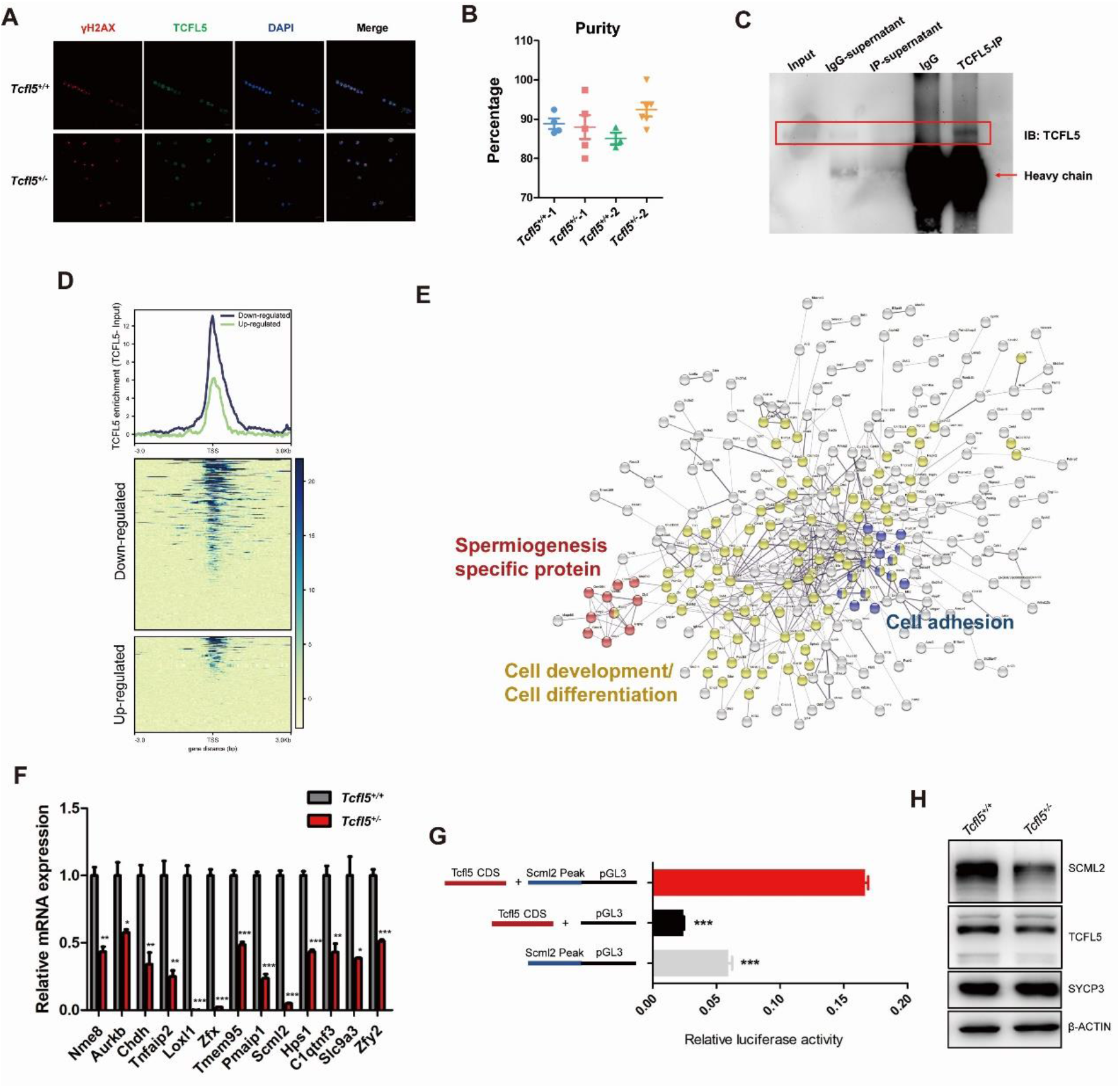
Analysis of genes transcriptionally regulated by TCFL5. (A) Validation of purified pachytene spermatocytes. The isolated cells from *Tcfl5*^+/+^ and *Tcfl5*^+/−^ mice were immunostained for γH2AX (red) and TCFL5 (green). γH2AX, a DNA-break marker restricted to the XY body was used to verify pachytene spermatocytes; DNA was stained with DAPI (scale bars, 20 μm). (B) Isolated cell purity is represented as mean ± SEM. (C) Western blots of TCFL5 protein to validate the enrichment of TCFL5 protein by ChIP experiments. (D) Distribution of TCFL5 ChIP reads on gene bodies from downregulated and upregulated genes, and heatmap of TCFL5-binding signals from −kb to +3 kb from the TSS of genes. (E) STRING protein network analysis of downregulated genes in pachytene spermatocytes from *Tcfl5*^+/−^ mice. (F) The expression validation by q-PCR of genes highly correlated with mouse spermatogenesis. (G) Dual-luciferase reporter assay confirming the promotion of peak sequence by *Scml2* (data are presented as mean ± s.d., n=3 per group, *** indicates P < 0.001 by unpaired Student’s *t test*). (K) Western blots indicate the diminished expression of SCML2 in *Tcfl5*^+/−^ compared to that in *Tcfl5*^+/+^ mouse testes. TCFL5, β-ACTIN, and SYCP3 served as controls.

**Fig. S3.**
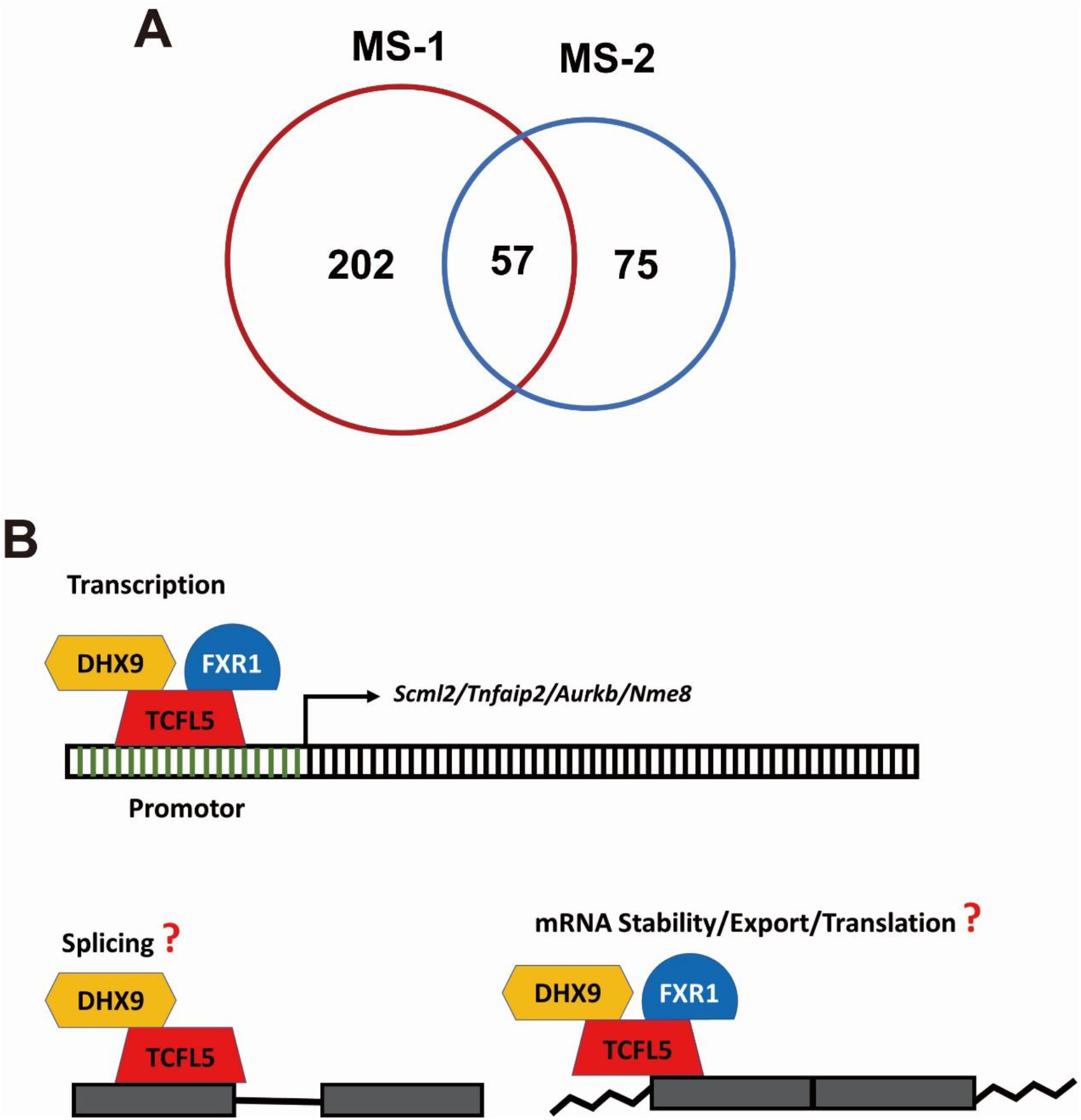
Proposed model of TCFL5-mediated regulatory mechanisms. (A) Protein overlap between two independent experiments of TCFL5 immunoprecipitation and protein mass spectrometry. (B) A proposed regulatory model of TCFL5 in the testes. TCFL5 transcriptionally regulates the expression of genes essential for spermatocyte and spermatid development—including *Scml2*, *Tnfaip2*, *Aurkb,* and *Nme8*. In addition, TCFL5 might interact with DHX9 and FXR1 to regulate mRNA fates such as splicing at posttranscriptional levels.

Table S1. List of genes differently expressed in pachytene spermatocytes between *Tcfl5*^+/+^ and *Tcfl5*^+/−^ mice.

Table S2. TCFL5-bound genomic loci in wild-type mouse testes at P21.

Table S3. Proteins identified in the mass spectrometry experiments using mouse testes at P21.

## Referneces

Adams, R.R., M. Carmena, and W.C. Earnshaw. (2001). Chromosomal passengers and the (aurora) ABCs of mitosis. Trends in cell biology 11, 49–54. doi:10.1016/s0962-8924(00)01880-8

Adams, S.R., S. Maezawa, K.G. Alavattam, H. Abe, A. Sakashita, M. Shroder, T.J. Broering, J. Sroga Rios, M.A. Thomas, X. Lin, C.M. Price, A. Barski, P.R. Andreassen, and S.H. Namekawa. (2018). RNF8 and SCML2 cooperate to regulate ubiquitination and H3K27 acetylation for escape gene activation on the sex chromosomes. PLoS Genet 14, e1007233. doi:10.1371/journal.pgen.1007233

Bai, S., and K. Fu. (2018). Sox30 initiates transcription of haploid genes during late meiosis and spermiogenesis in mouse testes. 145. doi:10.1242/dev.164855

Ballow, D., M.L. Meistrich, M. Matzuk, and A. Rajkovic. (2006a). Sohlh1 is essential for spermatogonial differentiation. Dev Biol 294, 161–167. doi:10.1016/j.ydbio.2006.02.027

Ballow, D.J., Y. Xin, Y. Choi, S.A. Pangas, and A. Rajkovic. (2006b). Sohlh2 is a germ cell-specific bHLH transcription factor. Gene Expr Patterns 6, 1014–1018. doi:10.1016/j.modgep.2006.04.007

Bao, J., C. Tang, J. Li, Y. Zhang, B.P. Bhetwal, H. Zheng, and W. Yan. (2014). RAN-binding protein 9 is involved in alternative splicing and is critical for male germ cell development and male fertility. PLoS Genet 10, e1004825. doi:10.1371/journal.pgen.1004825

Bettegowda, A., and M.F. Wilkinson. (2010). Transcription and post-transcriptional regulation of spermatogenesis. Philos. Trans. R. Soc. Lond. B Biol. Sci. 365, 1637–1651. doi:10.1098/rstb.2009.0196

Creyghton, M.P., A.W. Cheng, G.G. Welstead, T. Kooistra, B.W. Carey, E.J. Steine, J. Hanna, M.A. Lodato, G.M. Frampton, P.A. Sharp, L.A. Boyer, R.A. Young, and R. Jaenisch. (2010). Histone H3K27ac separates active from poised enhancers and predicts developmental state. Proc Natl Acad Sci U S A 107, 21931–21936. doi:10.1073/pnas.1016071107

Davidovic, L., N. Durand, O. Khalfallah, R. Tabet, P. Barbry, B. Mari, S. Sacconi, H. Moine, and B. Bardoni. (2013). A novel role for the RNA-binding protein FXR1P in myoblasts cell-cycle progression by modulating p21/Cdkn1a/Cip1/Waf1 mRNA stability. PLoS Genet 9, e1003367. doi:10.1371/journal.pgen.1003367

Davis, R.L., H. Weintraub, and A.B. Lassar. (1987). Expression of a single transfected cDNA converts fibroblasts to myoblasts. Cell 51, 987–1000.

Dictenberg, J.B., S.A. Swanger, L.N. Antar, R.H. Singer, and G.J. Bassell. (2008). A direct role for FMRP in activity-dependent dendritic mRNA transport links filopodial-spine morphogenesis to fragile X syndrome. Dev Cell 14, 926–939. doi:10.1016/j.devcel.2008.04.003

Eddy, E.M. (2002). Male germ cell gene expression. Recent Prog Horm Res 57, 103–128. doi:10.1210/rp.57.1.103

Fan, Y., J. Yue, M. Xiao, H. Han-Zhang, Y.V. Wang, C. Ma, Z. Deng, Y. Li, Y. Yu, X. Wang, S. Niu, Y. Hua, Z. Weng, P. Atadja, E. Li, and B. Xiang. (2017). FXR1 regulates transcription and is required for growth of human cancer cells with TP53/FXR2 homozygous deletion. Elife 6. doi:10.7554/eLife.26129

Filippova, G.N., S. Fagerlie, E.M. Klenova, C. Myers, Y. Dehner, G. Goodwin, P.E. Neiman, S.J. Collins, and V.V. Lobanenkov. (1996). An exceptionally conserved transcriptional repressor, CTCF, employs different combinations of zinc fingers to bind diverged promoter sequences of avian and mammalian c-myc oncogenes. Molecular and cellular biology 16, 2802–2813. doi:10.1128/mcb.16.6.2802

Geuens, T., D. Bouhy, and V. Timmerman. (2016). The hnRNP family: insights into their role in health and disease. Hum Genet 135, 851–867. doi:10.1007/s00439-016-1683-5

Hao, J., M. Yamamoto, T.E. Richardson, K.M. Chapman, B.S. Denard, R.E. Hammer, G.Q. Zhao, and F.K. Hamra. (2008). Sohlh2 knockout mice are male-sterile because of degeneration of differentiating type A spermatogonia. Stem Cells 26, 1587–1597. doi:10.1634/stemcells.2007-0502

Hartman, T.R., S. Qian, C. Bolinger, S. Fernandez, D.R. Schoenberg, and K. Boris-Lawrie. (2006). RNA helicase A is necessary for translation of selected messenger RNAs. Nat Struct Mol Biol 13, 509–516. doi:10.1038/nsmb1092

Hartmuth, K., H. Urlaub, H.P. Vornlocher, C.L. Will, M. Gentzel, M. Wilm, and R. Lührmann. (2002). Protein composition of human prespliceosomes isolated by a tobramycin affinity-selection method. Proc Natl Acad Sci U S A 99, 16719–16724. doi:10.1073/pnas.262483899

Hasegawa, K., H.S. Sin, S. Maezawa, T.J. Broering, A.V. Kartashov, K.G. Alavattam, Y. Ichijima, F. Zhang, W.C. Bacon, K.D. Greis, P.R. Andreassen, A. Barski, and S.H. Namekawa. (2015). SCML2 establishes the male germline epigenome through regulation of histone H2A ubiquitination. Dev Cell 32, 574–588. doi:10.1016/j.devcel.2015.01.014

Hou, L., Y. Wei, Y. Lin, X. Wang, Y. Lai, M. Yin, Y. Chen, X. Guo, S. Wu, Y. Zhu, J. Yuan, M. Tariq, N. Li, H. Sun, H. Wang, X. Zhang, J. Chen, X. Bao, and R. Jauch. (2020). Concurrent binding to DNA and RNA facilitates the pluripotency reprogramming activity of Sox2. Nucleic acids research 48, 3869–3887. doi:10.1093/nar/gkaa067

Hsu, S.H., H.M. Hsieh-Li, and H. Li. (2004). Dysfunctional spermatogenesis in transgenic mice overexpressing bHLH-Zip transcription factor, Spz1. Exp Cell Res 294, 185–198. doi:10.1016/j.yexcr.2003.10.029

Hudson, W.H., and E.A. Ortlund. (2014). The structure, function and evolution of proteins that bind DNA and RNA. Nature reviews. Molecular cell biology 15, 749–760. doi:10.1038/nrm3884

Jeon, Y., and J.T. Lee. (2011). YY1 tethers Xist RNA to the inactive X nucleation center. Cell 146, 119–133. doi:10.1016/j.cell.2011.06.026

Ji, X., Y. Zhou, S. Pandit, J. Huang, H. Li, C.Y. Lin, R. Xiao, C.B. Burge, and X.D. Fu. (2013). SR proteins collaborate with 7SK and promoter-associated nascent RNA to release paused polymerase. Cell 153, 855–868. doi:10.1016/j.cell.2013.04.028

Kashiwabara, S., J. Noguchi, T. Zhuang, K. Ohmura, A. Honda, S. Sugiura, K. Miyamoto, S. Takahashi, K. Inoue, A. Ogura, and T. Baba. (2002). Regulation of spermatogenesis by testis-specific, cytoplasmic poly(A) polymerase TPAP. Science (New York, N.Y.) 298, 1999–2002. doi:10.1126/science.1074632

Kashiwabara, S., T. Zhuang, K. Yamagata, J. Noguchi, A. Fukamizu, and T. Baba. (2000). Identification of a novel isoform of poly(A) polymerase, TPAP, specifically present in the cytoplasm of spermatogenic cells. Developmental biology 228, 106–115. doi:10.1006/dbio.2000.9894

Kimmins, S., C. Crosio, N. Kotaja, J. Hirayama, L. Monaco, C. Höög, M. van Duin, J.A. Gossen, and P. Sassone-Corsi. (2007). Differential functions of the Aurora-B and Aurora-C kinases in mammalian spermatogenesis. Molecular endocrinology (Baltimore, Md.) 21, 726–739. doi:10.1210/me.2006-0332

Kosir, R., P. Juvan, M. Perse, T. Budefeld, G. Majdic, M. Fink, P. Sassone-Corsi, and D. Rozman. (2012). Novel insights into the downstream pathways and targets controlled by transcription factors CREM in the testis. PLoS One 7, e31798. doi:10.1371/journal.pone.0031798

Kung, J.T., B. Kesner, J.Y. An, J.Y. Ahn, C. Cifuentes-Rojas, D. Colognori, Y. Jeon, A. Szanto, B.C. del Rosario, S.F. Pinter, J.A. Erwin, and J.T. Lee. (2015). Locus-specific targeting to the X chromosome revealed by the RNA interactome of CTCF. Molecular cell 57, 361–375. doi:10.1016/j.molcel.2014.12.006

Legrand, J.M.D., and R.M. Hobbs. (2018). RNA processing in the male germline: Mechanisms and implications for fertility. Semin Cell Dev Biol 79, 80–91. doi:10.1016/j.semcdb.2017.10.006

Lin, S., and X.D. Fu. (2007). SR proteins and related factors in alternative splicing. Advances in experimental medicine and biology 623, 107–122. doi:10.1007/978-0-387-77374-2_7

Martianov, I., M.A. Choukrallah, A. Krebs, T. Ye, S. Legras, E. Rijkers, W. Van Ijcken, B. Jost, P. Sassone-Corsi, and I. Davidson. (2010). Cell-specific occupancy of an extended repertoire of CREM and CREB binding loci in male germ cells. BMC Genomics 11, 530. doi:10.1186/1471-2164-11-530

Maruyama, O., H. Nishimori, T. Katagiri, Y. Miki, A. Ueno, and Y. Nakamura. (1998). Cloning of TCFL5 encoding a novel human basic helix-loop-helix motif protein that is specifically expressed in primary spermatocytes at the pachytene stage. Cytogenet Cell Genet 82, 41–45. doi:10.1159/000015061

Massari, M.E., and C. Murre. (2000). Helix-loop-helix proteins: regulators of transcription in eucaryotic organisms. Molecular and cellular biology 20, 429–440. doi:10.1128/mcb.20.2.429-440.2000

Matveeva, E., J. Maiorano, Q. Zhang, A.M. Eteleeb, and P. Convertini. (2016). Involvement of PARP1 in the regulation of alternative splicing. 2, 15046. doi:10.1038/celldisc.2015.46

Matzuk, M.M., and D.J. Lamb. (2008). The biology of infertility: research advances and clinical challenges. Nature medicine 14, 1197–1213. doi:10.1038/nm.f.1895

Melikishvili, M., J.H. Chariker, E.C. Rouchka, and Y.N. Fondufe-Mittendorf. (2017). Transcriptome-wide identification of the RNA-binding landscape of the chromatin-associated protein PARP1 reveals functions in RNA biogenesis. Cell discovery 3, 17043. doi:10.1038/celldisc.2017.43

Miranda-Vizuete, A., K. Tsang, Y. Yu, A. Jiménez, M. Pelto-Huikko, C.J. Flickinger, P. Sutovsky, and R. Oko. (2003). Cloning and developmental analysis of murid spermatid-specific thioredoxin-2 (SPTRX-2), a novel sperm fibrous sheath protein and autoantigen. The Journal of biological chemistry 278, 44874–44885. doi:10.1074/jbc.M305475200

Murre, C., P.S. McCaw, H. Vaessin, M. Caudy, L.Y. Jan, Y.N. Jan, C.V. Cabrera, J.N. Buskin, S.D. Hauschka, A.B. Lassar, and et al. (1989). Interactions between heterologous helix-loop-helix proteins generate complexes that bind specifically to a common DNA sequence. Cell 58, 537–544. doi:10.1016/0092-8674(89)90434-0

Nakajima, T., C. Uchida, S.F. Anderson, C.G. Lee, J. Hurwitz, J.D. Parvin, and M. Montminy. (1997). RNA helicase A mediates association of CBP with RNA polymerase II. Cell 90, 1107–1112. doi:10.1016/s0092-8674(00)80376-1

Nalam, R.L., and M.M. Matzuk. (2010). Local signalling environments and human male infertility: what we can learn from mouse models. Expert reviews in molecular medicine 12, e15. doi:10.1017/s1462399410001468

Nantel, F., L. Monaco, N.S. Foulkes, D. Masquilier, M. LeMeur, K. Henriksen, A. Dierich, M. Parvinen, and P. Sassone-Corsi. (1996). Spermiogenesis deficiency and germ-cell apoptosis in CREM-mutant mice. Nature 380, 159–162. doi:10.1038/380159a0

Olson, E.N. (1990). MyoD family: a paradigm for development? Genes & development 4, 1454–1461. doi:10.1101/gad.4.9.1454

Paronetto, M.P., V. Messina, E. Bianchi, M. Barchi, G. Vogel, C. Moretti, F. Palombi, M. Stefanini, R. Geremia, S. Richard, and C. Sette. (2009). Sam68 regulates translation of target mRNAs in male germ cells, necessary for mouse spermatogenesis. J Cell Biol 185, 235–249. doi:10.1083/jcb.200811138

Ranji, A., N. Shkriabai, M. Kvaratskhelia, K. Musier-Forsyth, and K. Boris-Lawrie. (2011). Features of double-stranded RNA-binding domains of RNA helicase A are necessary for selective recognition and translation of complex mRNAs. J Biol Chem 286, 5328–5337. doi:10.1074/jbc.M110.176339

Saldaña-Meyer, R., E. González-Buendía, G. Guerrero, V. Narendra, R. Bonasio, F. Recillas-Targa, and D. Reinberg. (2014). CTCF regulates the human p53 gene through direct interaction with its natural antisense transcript, Wrap53. Genes & development 28, 723–734. doi:10.1101/gad.236869.113

Schlegel, P.N. (2009). Evaluation of male infertility. Minerva ginecologica 61, 261–283.

Shi, Y., L. Zhang, S. Song, M.E. Teves, H. Li, Z. Wang, R.A. Hess, G. Jiang, and Z. Zhang. (2013). The mouse transcription factor-like 5 gene encodes a protein localized in the manchette and centriole of the elongating spermatid. Andrology 1, 431–439. doi:10.1111/j.2047-2927.2013.00069.x

Shukla, S., E. Kavak, M. Gregory, M. Imashimizu, B. Shutinoski, M. Kashlev, P. Oberdoerffer, R. Sandberg, and S. Oberdoerffer. (2011). CTCF-promoted RNA polymerase II pausing links DNA methylation to splicing. Nature 479, 74–79. doi:10.1038/nature10442

Siep, M., E. Sleddens-Linkels, S. Mulders, H. van Eenennaam, E. Wassenaar, W.A. Van Cappellen, J. Hoogerbrugge, J.A. Grootegoed, and W.M. Baarends. (2004). Basic helix-loop-helix transcription factor Tcfl5 interacts with the Calmegin gene promoter in mouse spermatogenesis. Nucleic Acids Res 32, 6425–6436. doi:10.1093/nar/gkh979

Sigova, A.A., B.J. Abraham, X. Ji, B. Molinie, N.M. Hannett, Y.E. Guo, M. Jangi, C.C. Giallourakis, P.A. Sharp, and R.A. Young. (2015). Transcription factor trapping by RNA in gene regulatory elements. Science 350, 978–981. doi:10.1126/science.aad3346

Silber, S.J. (1994). A modern view of male infertility. Reproduction, fertility, and development 6, 93–103; discussion 103-104. doi:10.1071/rd9940093

Smith, T.B., M.A. Baker, H.S. Connaughton, U. Habenicht, and R.J. Aitken. (2013). Functional deletion of Txndc2 and Txndc3 increases the susceptibility of spermatozoa to age-related oxidative stress. Free radical biology & medicine 65, 872–881. doi:10.1016/j.freeradbiomed.2013.05.021

Steigemann, P., C. Wurzenberger, M.H. Schmitz, M. Held, J. Guizetti, S. Maar, and D.W. Gerlich. (2009). Aurora B-mediated abscission checkpoint protects against tetraploidization. Cell 136, 473–484. doi:10.1016/j.cell.2008.12.020

Suzuki, H., H.W. Ahn, T. Chu, W. Bowden, K. Gassei, K. Orwig, and A. Rajkovic. (2012). SOHLH1 and SOHLH2 coordinate spermatogonial differentiation. Dev Biol 361, 301–312. doi:10.1016/j.ydbio.2011.10.027

Talasz, H., H.H. Lindner, B. Sarg, and W. Helliger. (2005). Histone H4-lysine 20 monomethylation is increased in promoter and coding regions of active genes and correlates with hyperacetylation. J Biol Chem 280, 38814–38822. doi:10.1074/jbc.M505563200

Toyoda, S., T. Miyazaki, S. Miyazaki, T. Yoshimura, M. Yamamoto, F. Tashiro, E. Yamato, and J. Miyazaki. (2009). Sohlh2 affects differentiation of KIT positive oocytes and spermatogonia. Dev Biol 325, 238–248. doi:10.1016/j.ydbio.2008.10.019

Vasudevan, S., and J.A. Steitz. (2007). AU-rich-element-mediated upregulation of translation by FXR1 and Argonaute 2. Cell 128, 1105–1118. doi:10.1016/j.cell.2007.01.038

Vasudevan, S., Y. Tong, and J.A. Steitz. (2007). Switching from repression to activation: microRNAs can up-regulate translation. Science (New York, N.Y.) 318, 1931–1934. doi:10.1126/science.1149460

Wang, Y., H. Wang, Y. Zhang, Z. Du, W. Si, S. Fan, D. Qin, M. Wang, Y. Duan, L. Li, Y. Jiao, Y. Li, Q. Wang, Q. Shi, X. Wu, and W. Xie. (2019). Reprogramming of Meiotic Chromatin Architecture during Spermatogenesis. Molecular cell 73, 547–561.e546. doi:10.1016/j.molcel.2018.11.019

Wolf, F.W., V. Sarma, M. Seldin, S. Drake, S.J. Suchard, H. Shao, K.S. O’Shea, and V.M. Dixit. (1994). B94, a primary response gene inducible by tumor necrosis factor-alpha, is expressed in developing hematopoietic tissues and the sperm acrosome. The Journal of biological chemistry 269, 3633–3640.

Xiao, R., J.Y. Chen, Z. Liang, D. Luo, G. Chen, Z.J. Lu, Y. Chen, B. Zhou, H. Li, X. Du, Y. Yang, M. San, X. Wei, W. Liu, E. Lécuyer, B.R. Graveley, G.W. Yeo, C.B. Burge, M.Q. Zhang, Y. Zhou, and X.D. Fu. (2019). Pervasive Chromatin-RNA Binding Protein Interactions Enable RNA-Based Regulation of Transcription. Cell 178, 107–121.e118. doi:10.1016/j.cell.2019.06.001

Zang, C., D.E. Schones, C. Zeng, K. Cui, K. Zhao, and W. Peng. (2009). A clustering approach for identification of enriched domains from histone modification ChIP-Seq data. Bioinformatics (Oxford, England) 25, 1952–1958. doi:10.1093/bioinformatics/btp340

Zhang, S., K. Buder, C. Burkhardt, B. Schlott, M. Görlach, and F. Grosse. (2002). Nuclear DNA helicase II/RNA helicase A binds to filamentous actin. J Biol Chem 277, 843–853. doi:10.1074/jbc.M109393200

Zhang, S., C. Herrmann, and F. Grosse. (1999). Pre-mRNA and mRNA binding of human nuclear DNA helicase II (RNA helicase A). J Cell Sci 112 (Pt 7), 1055–1064.

